# Filter paper disks as a matrix for manipulation of recombinant proteins

**DOI:** 10.1101/2022.03.23.485486

**Authors:** Eric H. Ball, Nicoletta T. Basilone

**Author notes:** **Corresponding Author** Eric H. Ball - Department of Biochemistry, University of Western Ontario, London, ON, Canada N6G3C3.

## Abstract

Filter paper provides an excellent matrix for retention of proteins containing a cellulose binding domain. To use this capability for manipulating recombinant fusion proteins, binding and elution parameters were explored and procedures developed for small scale purification, modification and assay. Proteins were tagged with the cellulose binding domain from the *C thermocellum* CipB gene via a cleavable linker. Filter paper disks of 6mm diameter were able to bind up to 80 μg protein although there was a substantial dependence on molecular size. Different means of introducing fusion proteins to the disks allow either binding within 20 minutes from microliter volumes or slower binding from milliliter volumes. Elution with protease in small volumes yielded greater than 10 μg amounts with concentrations in the 1-2 mg/ml range. To demonstrate their utility, disks were used for small scale protein purification, covalent modification of protein, immunoprecipitation, and in a binding assay. These versatile methods allow parallel processing of multiple samples and may find many uses when only small amounts of protein are needed.

## Introduction

Immobilization of proteins on a solid support is central to many biomolecular techniques, notably immunoprecipitations, affinity purifications and many binding assays. Thus a variety of methods have been developed to attach proteins to different types of surface, ranging from specific oriented attachment to the non-specific adhesion of proteins to nitrocellulose or plastic^1–3^. Covalent, permanent linkage can provide a high density affinity matrix for specific binding as in purification of IgG by protein A-sepharose^4^, and has the advantage that the attached protein does not elute even under denaturing conditions. Non-covalent binding to create affinity matrices has advantages in some circumstances, however; especially in ease of preparation and in allowing removal and analysis of the bound protein.

Filter paper has been a useful substrate for many reactions and assays (reviewed in Pelton, 2009^5^), particularly involving precipitation of proteins or nucleic acids. Filter paper is almost purely cellulose: it is hydrophilic with low background binding, easy to handle, commonly available and relatively cheap to buy. It has been used for the covalent attachment of peptides and proteins by several means including cyanogen bromide^6^ and other activating chemicals ^7,8^. A drawback, however, has been a relatively low coupling efficiency for proteins: on the order of a few micrograms to nanograms per square centimeter. This is adequate for some techniques, eg immunodetection, but much less so for others.

The discovery of cellulose binding domains from bacterial cellulosomes provided a useful affinity tag and led to the use of cellulose preparations for protein purifications^9^. These domains are a subset of carbohydrate-binding modules; a diverse set of proteins from many organisms, now divided into more than 80 families^9–12^. Some of these domains are relatively small, stable, bind cellulose with high affinity and can be expressed in *E coli*; excellent characteristics for a reagent for linking to filter paper or other forms of cellulose. Indeed, there are many examples of using CBMs to purify or immobilize fusion proteins^9,13–17^. In the present work we have investigated the parameters for binding, modification and elution of proteins fused with a cellulose binding domain from *Clostridium thermocellum*^18^ to a convenient size of filter paper disk in order to develop practical, useful methods for manipulating microgram amounts of proteins. These methods are likely to be useful in many situations, but particularly for the parallel study of small libraries of mutants when microgram quantities are sufficient for characterization.

## Materials and Methods

### Plasmid constructs

Proteins were expressed from modified pET vectors^19^. The backbone came from pET-21d (Novagen, Madison) modified to contain an N-terminal histag (from pET-14b (Novagen, Madison)) followed by an NdeI site for insertion of one coding sequence, followed by a TEV cleavage site (DIPTTENLYQSG), then a multiple cloning site for insertion of a second coding sequence.

The pEBC vector was produced by inserting the CBM coding sequence (amino acids 273-477 of cipB gene (Uniprot Q01866)) at the NdeI site. The DNA sequence was optimized for *E coli* expression and synthesized by IDT (Coralville, IA). The pEBT7 vector contained the trxA coding sequence (Uniprot P0AA25) at the NdeI site. The pEBCaM vector had calmodulin coding sequence (Uniprot P0DP23) at the NdeI site.

The following cDNA sequences of proteins or peptides were cloned into the pEBC vector to code for fusion proteins with CBM at the N-terminal: gfp (eGFP, uniprot C5MKY7); Vh (vinculin head region, amino acid 1-850 of human vinculin (Uniprot P18206)); VBS (vinculin binding sequence (amino acids 492-509 from invasin IpaA sequence (Uniprot P18010) containing an E494C mutation); Protein A (amino acids 154-269 from *S. aureus* protein A (Uniprot P02976)); human NQO1 (uniprot P15559); human NQO2 (uniprot P16083). NQO1 and NQO2 sequences were a gift from Dr. Brian Shilton^20,21^.

The calmodulin-GFP fusion was made by inserting the eGFP sequence into the pEBCaM vector multiple cloning site. The thioredoxin-Vh-GFP sequence was made by inserting the Vh sequence followed by the eGFP sequence into pEBT7. To make CBM with a calmodulin binding peptide, a designed calmodulin-binding peptide (sequence: ARWRNTIIAVTAANRFGN) was placed N-terminal to the CBM in pEBC and no TEV site was present.

### Protein purification

Several fusion proteins were purified by chromatography on Ni-NTA columns as follows: BL21 cells containing the appropriate plasmids were grown overnight at 37° in 200 ml of M9 medium containing 0.4% glucose, 0.2% casamino acids and 100 μg/ml ampicillin. The culture was diluted with 1.8 l of LB medium and shaken 2 h at 37°. The temperature was then lowered to 22.5°, IPTG added to 0.2 mM and incubation continued for 5 h. Cells were harvested by centrifugation, resuspended in 20 ml 10 mM Tris pH 7.5, 1 mM EDTA and frozen at -80° until needed. Cells were thawed, lysed by passage through an emulsiflex homogenizer (Avestin, Ottawa, Canada) at 15000 psi, NaCl was added to 0.3M, imidazole to 10 mM and the lysate centrifuged 1 h at 100,000 x g. The supernatant was applied to a column (∼6 ml) of Ni-NTA resin in an equilibration buffer of 10 mM Tris pH 7.5, 10 mM imidazole, 0.3 M NaCl, 0.03% (v/v) mercaptoethanol. After washing with equilibration buffer, protein was eluted with equilibration buffer containing 0.25 M imidazole.

Some proteins were subject to further purification. CaM-CBM and CaM_GFP were further purified by binding to a phenyl sepharose column, washing with 10 mM Tris pH 7.5, 0.5 mM CaCl_2_, 100 mM NaCl, then eluting with 10 mM Tris pH 7.5, 1 mM EDTA, 100 mM NaCl (TEN100 buffer).

TEV was expressed as the self-cleaving MBP fusion^22^ and purified on a Ni-NTA column as above.

### Use of filter paper disks

Disks were cut from Whatman #1 filter paper sheets using a 6 mm one hole punch and manipulated with flat tip tweezers. For protein binding, two protocols were used: 1) for long term immersion, disks were placed in a protein solution (in TEN100 buffer) in microcentrifuge tubes (1 ml volume) or 6 well tissue culture plates (2 ml volume) with rotation overnight at 4°C; 2) for short term binding, disks were placed on parafilm and protein solution directly pipetted on to them at room temperature. To prevent drying, wet disks were overlaid with parafilm and a thin glass sheet.

Following the protein binding step, disks were washed by placing individually on a sintered glass filter support base (2 cm diameter) with suction and 200 μl of buffer sucked through the disk.

To elute all the protein bound to a disk, it was immersed in 100 μl of 10 mM Tris pH 7.5, 1 mM EDTA, 1% SDS in a microcentrifuge tube and placed in a boiling water bath for 3 min.

To elute only the fusion partner of the CBM, disks were placed inside a small (0.2 ml) tube that had a small hole at the bottom (punctured with a hot needle). This tube was then placed inside a larger (0.5 ml) microcentrifuge tube and 7 μl of a solution of 100 μg/ml TEV in 10 mM Tris pH 7.5, 1 mM EDTA, 5 mM DTT pipetted on to the disk. Up to four disks could be done in one tube using 7 μl solution per disk. After a suitable length of time (usually 1 h at room temperature) the tubes were centrifuged for 3 sec. A further 7 μl of TEN-100 buffer per disk was pipetted directly on to the disk(s) followed by a second brief centrifugation.

### Small scale protein expression and isolation

A colony of BL21 pLysS transformed with fusion protein plasmid was picked and grown overnight at 37° in 2.5 ml terrific broth containing 100 μg/ml ampicillin, 34 μg/ml chloramphenicol and 0.2 mM IPTG. A 1.5 ml aliquot of this culture was centrifuged, resuspended in 0.75 ml of 10 mM Tris pH 7.5, 1 mM EDTA, subjected to two cycles of freeze/thaw, sonicated briefly (5 s) to decrease viscosity, and centrifuged 15 min at 20,000 x g in a microfuge. The supernatant was incubated with disks rotating overnight at 4° for protein binding.

### Enzyme activity assay

Enzyme was added to 1 ml 0.1 M Tris HCl, pH 7.5, 50 μM menadione, containing either 160 μM NADH or 140 μM BNAH. Absorbance at 340 nm (NADH) or 355 nm (BNAH) was recorded over several minutes. Enzyme amounts were adjusted to give a linear output over 2 minutes. The slope of the line was converted to concentration change/sec using extinction coefficients of 6.22 mM^-1^cm^-1^ for NADH and 7.24 mM^-1^cm^-1^ for BNAH.

### Immunoprecipitation

For disk-based immunoprecipitation, IgG (preimmune or anti-thioredoxin, Sigma T-0803) was mixed with SPA-CBM to concentrations of 4.5 mg/ml and 1.5 mg/ml respectively, and 4.5 ul was spotted directly on a filter paper disk. After 20 minutes under a parafilm cover the disks were washed, then rotated with 0.5 ml of *E coli* cell extract containing thioredoxin-Vh-GFP protein overnight at 4°. Disks were washed with 200 ul TEN100 buffer, then eluted with TEV treatment.

### Fluorescent labelling

Three disks were treated with the partially purified VBS-CBM 150 μg/ml in 2 ml TEN100 buffer containing 0.1 mM TCEP overnight at 4°. After washing with suction, 4.5 ul of a 1 mg/ml solution of fluorescein maleimide in buffer was pipetted onto the disk. After 1 hour incubation at room temperature in the dark under parafilm, the disks were washed with 200 μl buffer, then eluted as above either with SDS buffer or TEV.

### Calmodulin binding assay

CBP-CBM was bound to disks by spotting 4.5 ul of buffer containing 0.2 μg protein directly on each disk, incubating under parafilm for 20 min, then washing with 200 μl of binding buffer (10 mM Tris-HCl pH 7.5, 100 mM NaCl, 100 μg/ml casein, 0.5 mM CaCl_2_). Triplicate disks were incubated in 2 ml binding buffer with various concentrations of calmodulin-GFP overnight at 4°. Disks were then washed, placed on parafilm, overlaid with another layer of parafilm and a glass plate, and photographed with a BioRad ChemiDoc MP system using settings for GFP. For quantitation, spot densitometry with ImageLab software was performed on the image.

### Other methods

Total protein was measured by a modified Lowry procedure after TCA precipitation^23^. SDS PAGE used the buffer system of Laemmli^24^. Fluorescent images were taken with a BioRad ChemiDoc MP system using settings for GFP.

## Results

Filter paper is a convenient, easily manipulated source of cellulose for CBM domain binding. Out of several shapes and sizes tested, the most useful for fitting into tubes and that was easily obtained was a 6 mm diameter circle punched out of a Whatman No. 1 filter paper sheet. These disks weighed 2.85 ± 0.1 mg each, and could absorb 4 μl of water. Following wetting and brief drying under suction 2.8 μl of liquid was retained whereas after a further short centrifugation (3 sec) 1.9 μl remained. Disks were readily grasped with thin, flat-tipped tweezers (Dumont type 2a was particularly appropriate) and could be bent to fit into 0.2 ml tubes for centrifugation.

In order to develop procedures to use the disks for protein immobilization, a number of fusion proteins were produced and partially purified. The *C thermocellum* CBM is known to have a stable folded structure and bind cellulose with a Kd of ∼0.1 μM^9,18^. It proved to be a robust fusion partner for all proteins/domains that were tested. Figure 1 shows an SDS PAGE of the various partially purified proteins. Some contaminating bands remained, notably in the VBS (lane 4) and higher molecular weight Vh (lane 7) fusions. With the exception of the VBS, good yields of recombinant protein were obtained, ranging from 15 to 60 milligrams/liter of culture. The VBS fusion eluted from a nickel column was highly contaminated with a 70 kDa protein (Fig. 1, lane 4), possibly a chaperone, nevertheless the CBM was still active and able to bind cellulose.

**Figure 1.**
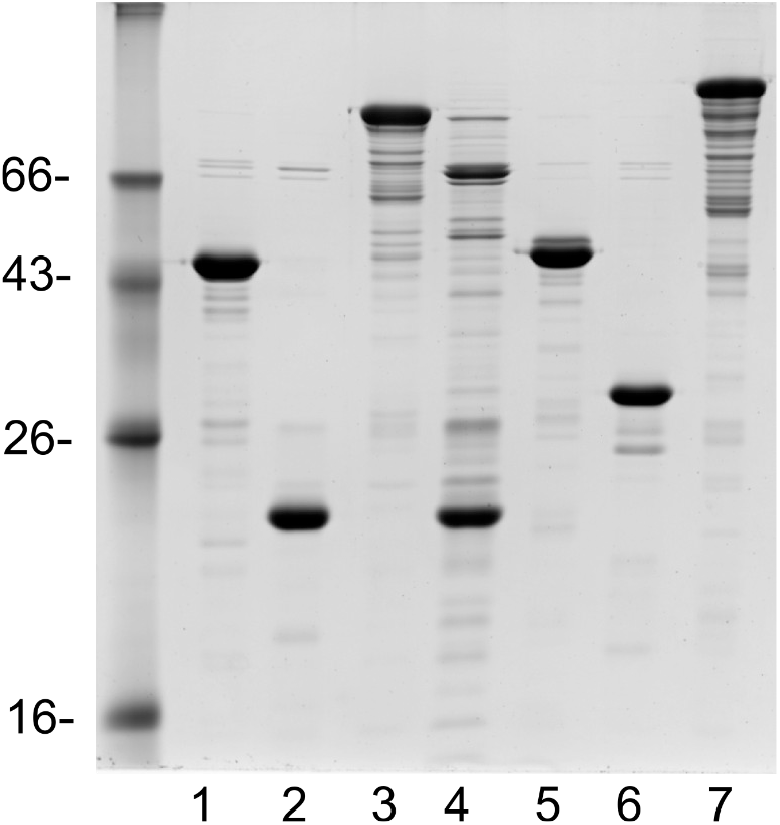
12% SDS PAGE of protein preparations used. Two micrograms of each were loaded and the gel was stained with Coomassie blue. Proteins are: lane 1: GFP-CBM (expected Mr 49.2 kDa), lane 2 CBP-CBM (22.7 kDa), lane 3: Vh-CBM (115 kDa), lane 4 E3C-CBM (24.8 kDa), lane 5 CaM-GFP (48 kDa), lane 6 Protein A-CBM (35 kDa), lane 7 Trx-GFP-Vh 136 kDa). Different degrees of purification obtained with different proteins.

### Binding of fusion proteins to disks

Removal of unbound proteins is important for many uses. To determine the effectiveness of washing for clearing of unbound protein from disks, 4 μl of a concentrated solution of BSA was pipetted directly onto disks, followed by washing with buffer. A 200 μl buffer wash under vacuum with left 2.7 ± 0.3% of the BSA on the disk, two successive washes of 200 μl each left 1.46±0.02%, and two washes separated by a 10 minute incubation in buffer left 0.41±0.02%. Thus a single wash removes 97% of unbound protein: this was sufficient for most purposes.

A high specific binding capacity would make the disks more useful. Binding of CBM fusion proteins to the disks was first examined by incubating disks in 2 ml of protein solution in a 6 well plate at 4°C. Different sizes of protein were used to look for effects of molecular weight. This protocol resulted in extensive binding to the disks (Fig. 2) with 82 μg of CBP-CBM protein/disk after 48 hours. Two phases of uptake were apparent: a faster initial phase over 5 hours followed by a slower binding that continued for at least 48 h. The capacity of the disks varied with the size of the fusion partner, with much greater binding by the CBP-CBM fusion (23 kDa) than GFP-CBM (49 kDa) and the Vh-CBM (115 kDa) although a significant amount was bound in all cases (30 - 80 μg) with very little non-specific binding. The dependence on size could be due to diffusion limitations or steric hindrance of closely spaced binding sites on cellulose.

**Figure 2.**
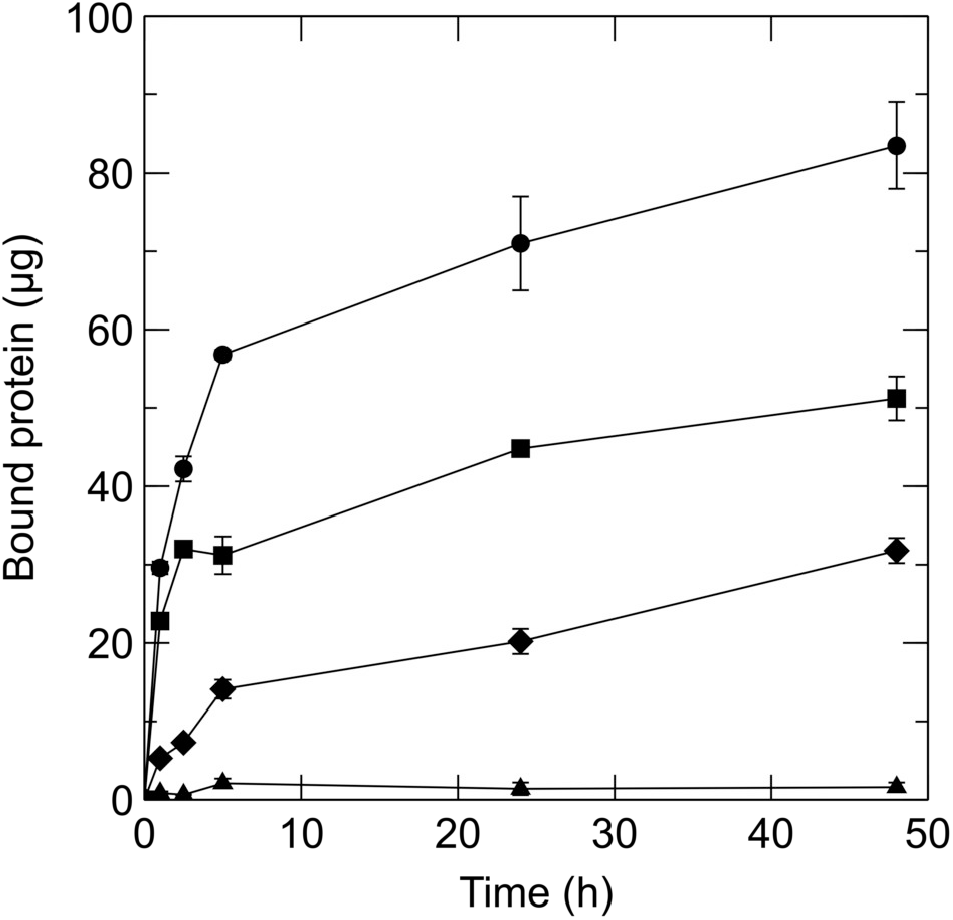
Protein binding to disks from solution. Disks were incubated in a 1.5 mg/ml solution of CBP-CBM (●), GFP-CBM (■), Vh-CBM (♦) or BSA (▲). Total protein bound to disks was determined after washing and elution in SDS solution. Triplicates ±SEM.

Binding was much faster using a direct pipetting protocol, wherein 4.5 μl of a protein solution was added to each dry disk, then covered with parafilm to prevent evaporation. Under these conditions, with a more concentrated protein solution, binding reached near maximal in five minutes (Fig. 3, left panel). While the kinetics of binding were similar for different protein concentrations, the final amount bound increased at higher input concentration. At 34 mg/ml GFP-CBM maximal capacity was ∼42 ug, corresponding to a 0.2 mM concentration within the disk, similar to the amount of binding observed after 24 hours incubation with 0.75 mg/ml protein in 2 ml of solution (Fig. 2). The direct spotting protocol provided a faster, convenient way to get proteins bound. Desorption from the disk after binding and washing was fairly slow: in three experiments with GFP-CBM the desorption after 24 hours soaking in buffer averaged 4% of the initial amount bound.

**Figure 3.**
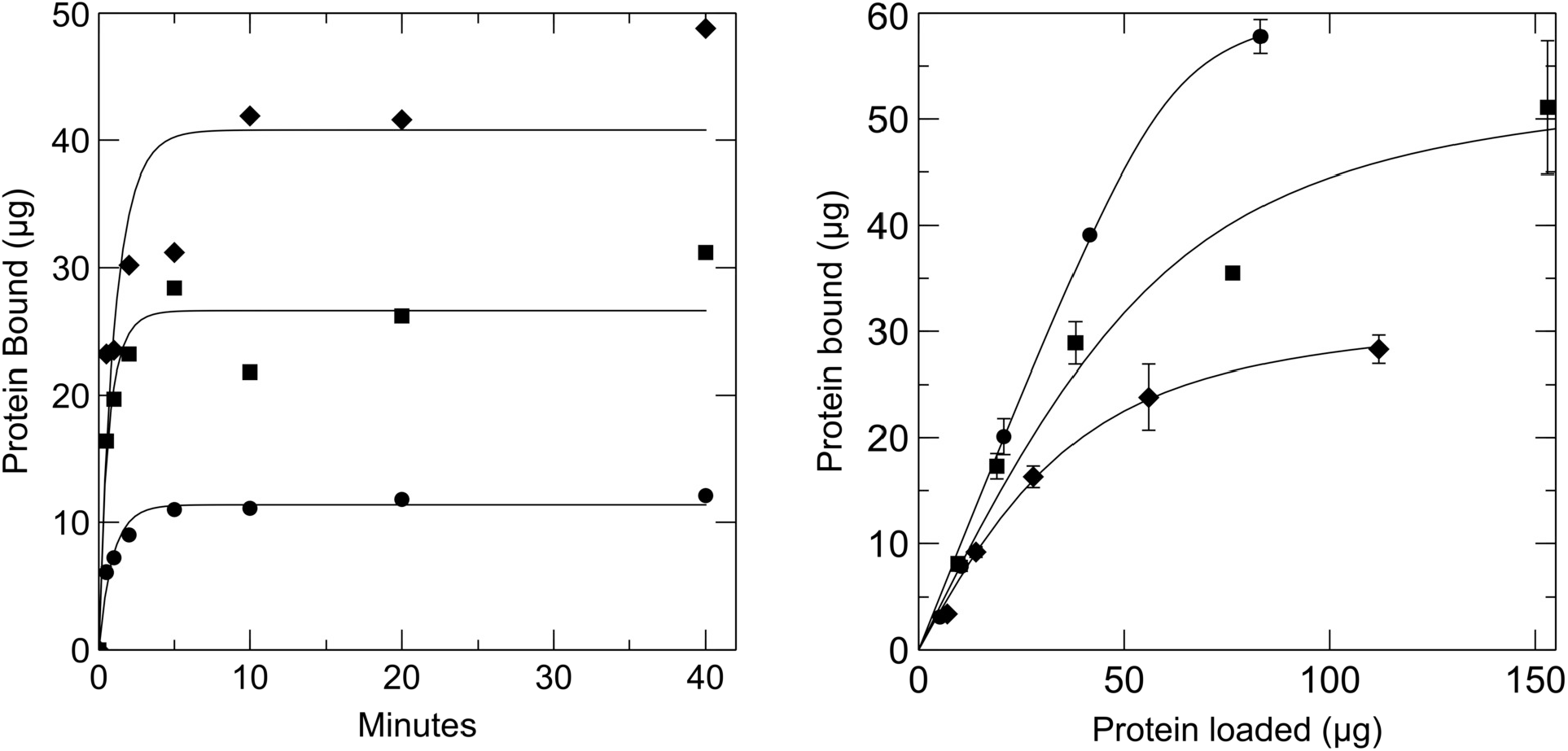
Binding to disks after direct application. Left panel: Different concentrations of GFP-CBM were spotted on triplicate disks for various times and bound protein determined after washing. Concentrations were 2.7 mg/ml (●), 7.2 mg/ml (■), and 34 mg/ml (♦). Progress curves were fitted to the data using the dynafit program^25^. Right panel: Triplicate disks were treated with various amounts of protein in 4.5 μl of solution for 20 min and bound protein subsequently measured. Fusion proteins were: CBP-CBM (●), GFP-CBM (■) and Vh-CBM (♦). Data are shown as average ± SEM fitted to an equilibrium binding model.

Using the direct spotting method, the amount of protein bound was investigated as a function of the amount initially added to the disk (Fig. 3, right panel). The amount bound increased with loading in a nearly linear fashion at lower amounts but followed a saturation curve visible at higher levels. Again, the effect of protein size in limiting binding was evident. In terms of molar amounts, 10 times as much CBP-CBM was bound per disk (2.5 nmol) as Vh-CBM (0.22 nmol) at the highest concentrations used. If a highly concentrated solution of protein is available, then this method is much quicker, but loading from more dilute solution requires the longer incubation in larger volume.

To specifically remove the fusion partner of the CBM, TEV was used to cleave in the linker region of the different fusion proteins. TEV was active in the disk (Fig. 4) and a 0.1 mg/ml concentration cut most of the fusion protein within 30 minutes although digestion continued for over 2 hours. Seven microliters of TEV solution per disk was used to allow for some evaporation over this time period. Based on these results, a one hour treatment time at room temperature was adopted as standard. Under these conditions 15±1.3 μg of GFP in 14 μl was recovered per disk; a yield of about 65% of theoretical.

**Figure 4.**
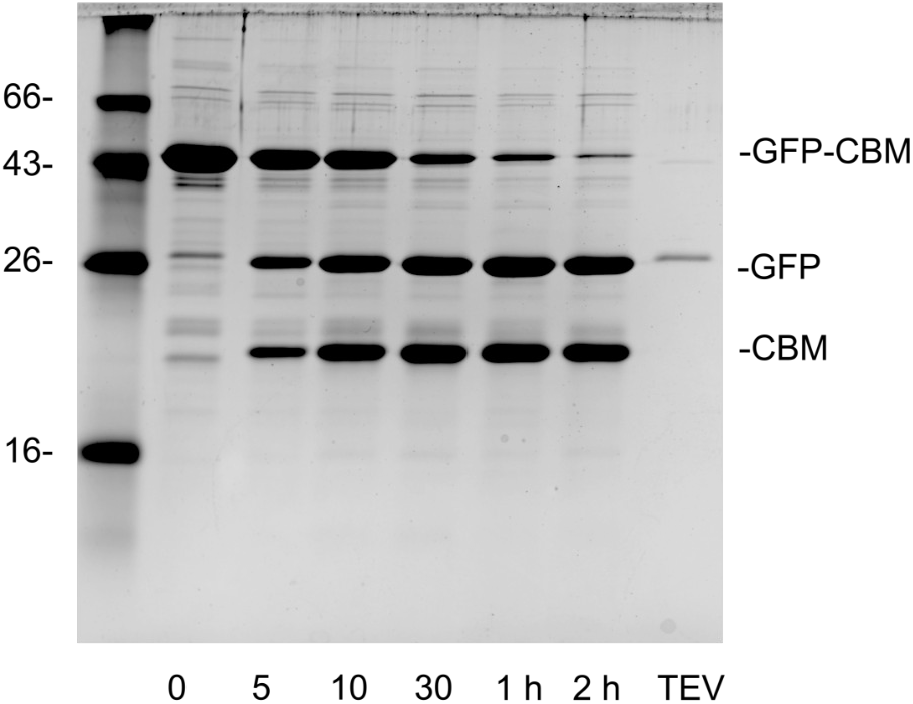
Time course of TEV digest of GFP-CBM bound to disk. Disks containing bound GFP-CBM were washed, placed in a 0.2 ml tube, centrifuged briefly and treated with 7 μl of 0.125 mg/ml TEV for various times indicated (minutes). The disk was then boiled in an SDS solution and one tenth of the solution loaded. Right lane shows TEV only.

### Applications of disk binding method

With some of the basic parameters evaluated, several applications became feasible. One possible use of the disk procedure is simply to isolate microgram quantities of a protein directly from a cell lysate. As a test of this method, human NQO1 (NAD(P)H:quinone oxidoreductase 1) and NQO2 enzymes were used. These enzymes are dimeric flavoproteins that nonetheless fold properly in E coli but differ from each other in their substrate specificity for NADH^26^. The cDNA sequences were placed in pEBC plasmid to make the CBM fusion, then expressed in BL21 pLysS E coli cells in small overnight cultures. The pLysS plasmid allowed a simple means to break the cells by freeze-thaw^19^. After lysis, supernatants from the centrifuged lysates were placed in a 6-well tissue culture plate, 4 disks added to each well and rotated overnight at 4°C. As shown in Figure 5, the disks acted as an excellent matrix for obtaining the enzymes in highly purified form (lanes 4, 6) from the cell extracts (odd lanes). Non-specific background (lane 2) was minimal. Total yields of the enzymes were 46 and 70 μg; whereas the activity measurements required 50 - 250 ng/assay. Reduction of menadione, a quinone stubstrate for the enzymes, measured with NADH as co-substrate gave values of 78±3 s^-1^ and 1.4±0.3 s^-1^ for NQO1 and NQO2, respectively, demonstrating the much higher efficacy of NQO1, as expected with NADH. The blank eluate showed no activity with either NADH or BNAH. On the other hand both enzymes had relatively high activities when the artificial co-substrate BNAH was used (348±34 s^-1^ and 101±6 s^-1^). The difference between the two enzymes use of NADH is a puzzling characteristic^27^ (reviewed in Vella et al^28^) and suggests NQO2 may function as a signalling protein rather than in detoxifying^20^.

**Figure 5.**
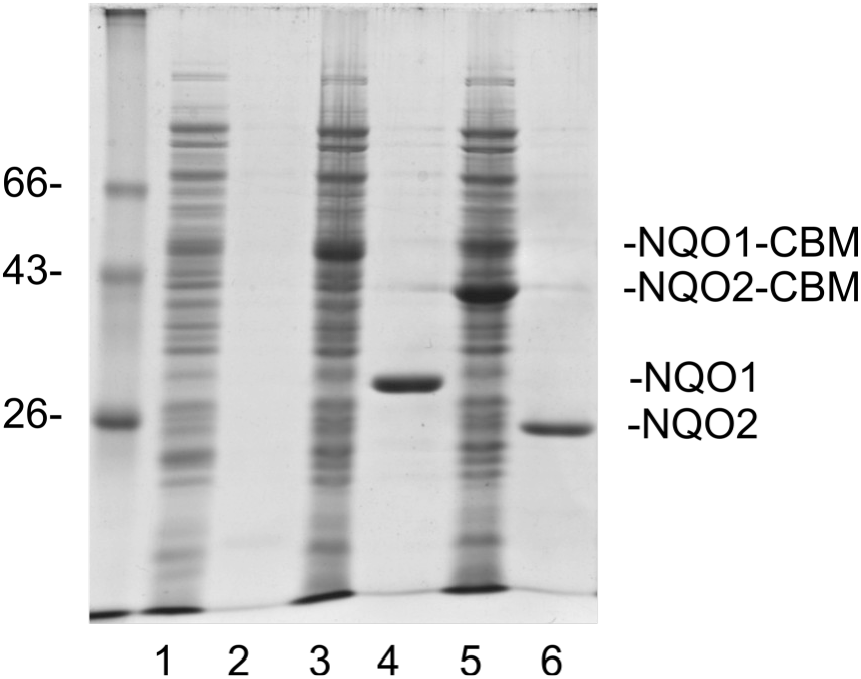
SDS PAGE showing purification of NQO2 enzymes using disks. Each pair of lanes shows cell extract followed by eluted protein. Lanes 1, 2; CBP-CBM (control), lanes 3, 4; NQO1-CBM, lanes 5, 6; NQO2-CBM.

Immobilizing proteins to a solid support can also provide a reaction chamber for covalent modification that allows easy removal of reagents. To test the disks for this purpose, a protein with a single exposed cysteine was made. A short peptide sequence (VBS) encoding 15 amino acids with a single cysteine residue was cloned into the pEBC vector and protein expressed. The CBM itself contains a single cysteine residue but it appears buried in the 3D structure^29^ and indeed did not react. After binding the fusion protein to disks, they were washed with TEN-100 buffer containing 0.1 mM TCEP to ensure reduction of cysteine. A solution of fluorescein maleimide was pipetted onto the disks and allowed to react for 1 h at room temperature before washing. Figure 6, lanes 1 show the fusion protein on the disk immediately after reaction (left panel; Coomassie stained, right panel; fluorescence). The right panel is a picture taken with fluorescent optics prior to staining the gel and shows that the fusion protein band has attached fluorescein. After TEV treatment (lanes 2) the free peptide and CBM appear. The CBM itself is non-fluorescent. The eluate from the disk (lanes 4) shows the now fluorescent peptide with some residual TEV visible on the stained gel (left panel). The CBM remains with the disk. Labelled proteins made in this manner could be used in a number of fluorescent assays.

**Figure 6.**
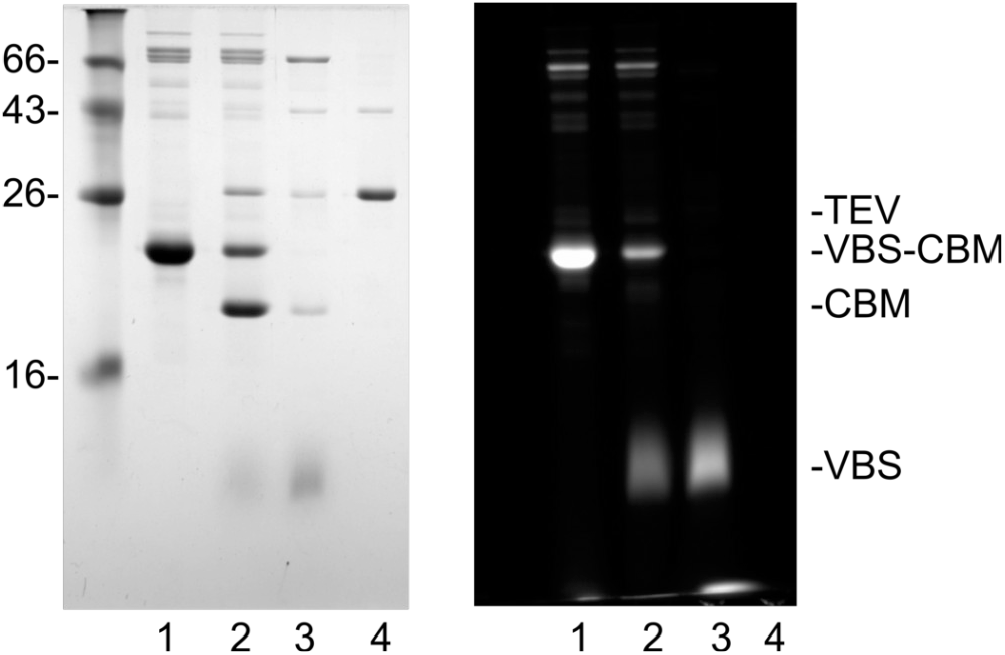
SDS PAGE showing cysteine modification of VBS peptide. VBS-CBM fusion protein was bound to disks overnight before washing and reaction with fluorescein-maleimide. Right panel shows a picture of the SDS gel taken with fluorescent optics before staining, and the left panel shows the gel after staining with Coomassie Blue. Lane 1: protein on disk after reaction, lane 2: protein on disk after TEV treatment, lane 3: protein eluted from disk, lane 4: TEV solution.

Procedures for immunoprecipitation generally involve using antibodies coupled or bound to a solid support. Indeed a CBM-Staphylococcal protein A (SPA-CBM) fusion has been previously used in an immunoassay^30,31^ and to purify IgG on cellulose substrates. To determine whether the filter paper disks could serve as an appropriate tool in immunoprecipitations, we constructed a similar SPA-CBM fusion. The fusion contained two IgG-binding repeats and was expressed and purified in high yields (see Fig 1, lane 6). The kinetics of binding soluble IgG to SPA-CBM disks was quite slow; instead mixing soluble SPA-CBM with IgG to create a complex before spotting on to dry disks gave more antibody incorporation into the disks. To demonstrate the use of the disks, anti-thioredoxin or nonimmune IgG with SPA-CBM was bound to disks and used to precipitate a thioredoxin-Vh-GFP fusion protein from a bacterial cell extract (Fig. 7). The loaded disks were incubated with a diluted cell extract overnight at 4° before washing and elution with TEV treatment. Lane 1 shows the elution from disks incubated without cell extract. The IgG heavy chain band is evident, whereas the light chains were visible as a diffuse band below the 26 kDa marker. When non-immune IgG was used with cell extract, the same bands are evident (lane 2) along with a few faint background bands. When the anti-thioredoxin IgG disk was incubated in the cell extract and eluted, a clear band of GFP-Vh is evident (lane 3). (The thioredoxin fusion protein has a TEV cleavage site, as does the SPA fusion, thus GFP-Vh is eluted along with the IgG.) Thus the disks with the SPA fusion protein have potential as an immunosorbent.

**Figure 7.**
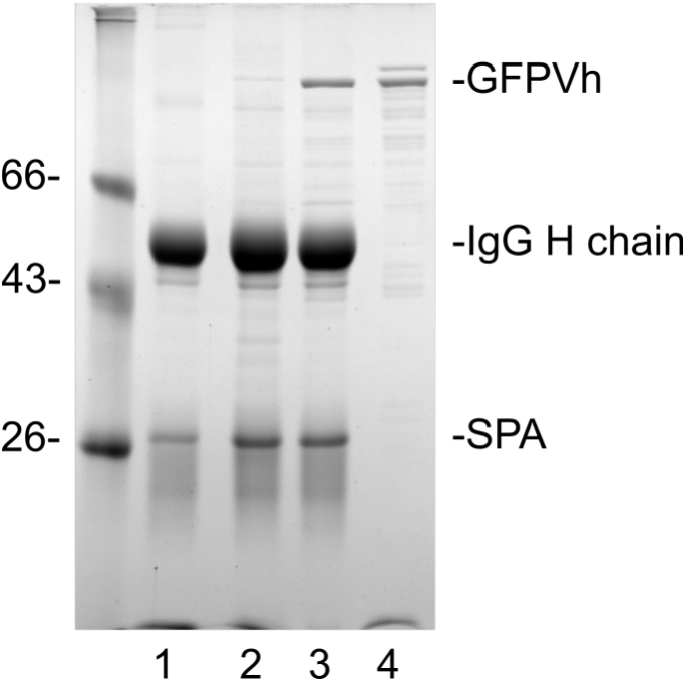
SDS PAGE showing immunoprecipitation using SPA-CBM fusion protein. SPA-CBM with IgG was spotted on a filter paper disk before incubation with a cell extract containing Trx-GFP-Vh fusion protein. Disks were treated with TEV and an aliquot of the eluate run on a 12% gel. Lane 1: anti-Trx IgG without cell extract, lane 2: non-immune IgG with cell extract, lane 3: anti-Trx IgG with cell extract, lane 4: purified GFP-Vh.

Immobilized proteins can provide a good substrate for binding assays, as notably in immunoassays with proteins bound to wells in plates^32^. The relatively quick washing procedure and the ability to read fluorescent signals directly off filter paper^33^ suggested that the protein bound disks might also be useful in binding assays. As a test binding assay the calmodulin - calmodulin-binding peptide (CBP) system was used. A fusion of a CBP to the CBM was bound to the disks, which were then incubated with various concentrations of a calmodulin-GFP fusion protein. After washing, fluorescence was quantitated on the disks. Figure 8 shows that a good fit to a hyperbolic binding curve was obtained. Curve fitting gave Kd of 4 nM for this interaction which is in the expected range, however, this must be regarded as an approximation given the possible problems of diffusion limitation, desorption of fusion protein, and the non-ideal distribution of binding sites (ie. in the disks rather than free in solution). Nonetheless this type of assay could certainly be used for comparative purposes or as a first step in screening for binding properties.

**Figure 8.**
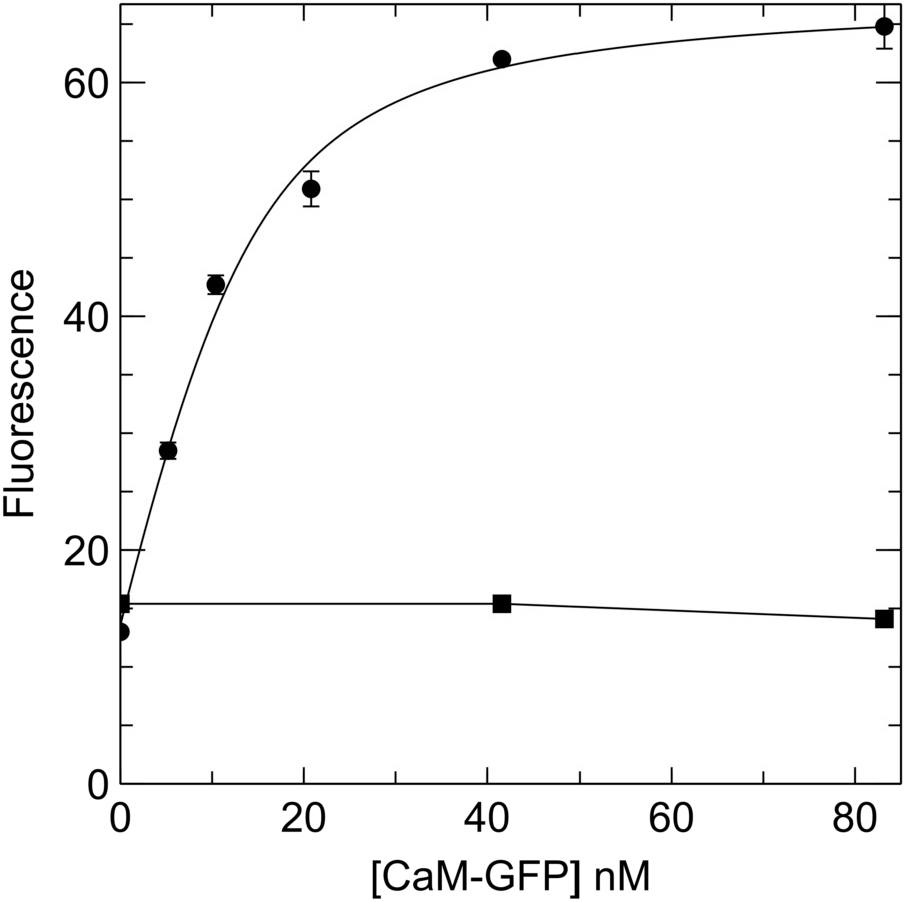
CaM-GFP binding to CBP-CBM on disks. Triplicate disks loaded with 0.2 μg each CBP-CBM were incubated with CaM-GFP in buffer containing 0.5 mM CaCl_2_ (●) or 1 mM EDTA (■) overnight at 4° C. Disks were washed, placed between layers of parafilm, imaged, and the intensity quantitated using Imagelab software. Points were fitted to a binding isotherm using Dynafit. Bars represent SEM.

## Discussion

Several cellulose-binding CBM domains have been used to tag many proteins for purification or immobilization (reviewed in Tomme et al^9^), proving to be well expressed and stable in *E coli* and yeast. The domain from *C thermocellum* cellulase is among the best characterized^29^, has relatively high affinity for cellulose, and can bind in high concentrations of urea^34^. It has been used to purify several fusion partners^30,35–37^ and for immunoassays ^38^. Various cellulosic substrates have been used with CBMs for binding or purification, including crystalline, amorphous, and fibers, but generally requiring columns or centrifugation to separate bound from unbound protein. To exploit these properties for easily manipulating small quantities of recombinant protein, new methods and a more convenient and ubiquitous substrate were required.

In developing these techniques, parameters for the binding and elution of fusion proteins to a convenient size of filter paper disk were explored and some possible specific uses tested. The size dependence of capacity (ranging from 0.26 nmol/disk for a 115 kDa protein to 3.5 nmol/disk for a 23 kDa protein) and long uptake times suggest slow diffusion to some sites within the disk. Likely internal sites within the disk are less accessible. The disks appear to have properties intermediate between crystalline and amorphous cellulose with regard to protein binding. The capacity of the disks appears greater than that of crystalline cellulose (∼8 μg/mg cellulose), but less than of amorphous cellulose (∼365 μg/mg^18^). The disks are considerably easier to work with, however^17^. When high concentrations of CBM fusion protein are available, the direct spotting technique allows for the fast loading of protein onto the disk. With lower concentrations, incubation for longer periods is needed for maximal binding.

When small amounts (<50 μg) of protein are needed, these disks have many advantages. Among them are the ease of use, ready availability, and economy of the cellulose substrate. Notably yields were sufficient for multiple enzyme assays of NQO1 and NQO2. Also notable are the speed of washing, elution in a small volume, oriented attachment of the fusion proteins, and the ability to elute bound proteins with SDS under denaturing conditions to check amount on disks or degradation. The ability to measure fluorescence directly from the disks opens up more possible uses for various types of binding assay. Because only small cultures are needed to produce enough fusion protein, multiple cultures can be processed in parallel, likely up to 50 or 100 at a time.

There are, however, a number of limitations that need to be considered. First, the protein must express and fold properly. Many eucaryotic proteins do not express well in *E coli*, but it is worth noting that the *C thermocellum* CBM has been expressed and used in yeast ^16,17^ and thus could also be applied to the paper disk technology described here. Second, the capacity for binding decreases with increasing size of the protein, thus protein complexes or larger proteins may not bind in usable amounts. However, the immunoprecipitation experiment shown in Figure 7 at least demonstrates that detectable amounts of large complexes (comprising SPA-CBM, IgG and a 136 kDa protein) will bind to the disks. Third, elution with TEV will leave the purified protein contaminated with a small amount of the protease, which could be problematic depending on the intended use. It would be possible to remove the protease using nickel beads, but more manipulation would be required. Finally, some desorption is likely to occur over longer time periods, if these are required.

The different applications exemplified here show that use of filter paper disks is a versatile technique that may be generally useful. When multiple mutants must be characterized (as in alanine scanning or cysteine scanning approaches, for examples) the parallel processing afforded by growing many small cultures and obtaining purified proteins simultaneously via this filter paper disk method will be very convenient. Likewise for deletion series, or random mutants at a single amino acid position in protein engineering applications. We are currently using the methods to examine multiple ancestral forms of the enzyme NQO2. No doubt many applications will be developed for specific tasks in the future.

## AUTHOR INFORMATION

### Author Contributions

EHB designed the project, performed experiments, analyzed data and wrote the manuscript. NTB performed experiments, helped write the manuscript and analyzed data.

### Notes

The authors declare no competing financial interests.

## ACKNOWLEDGMENTS

We thank Brian Shilton and Faiza Islam for helpful discussions and providing NQO1 and NQO2 DNA samples.

## ABBREVIATIONS

CBM: carbohydrate-binding module
TEV: tobacco etch virus protease
CaM: calmodulin
GFP: enhanced green fluorescent protein
SPA: *Staphylococcus aureus* protein A
NQO1: NAD(P)H:quinone oxidoreductase 1
BNAH: 1-benzyl-1,4-dihydro-3-pyridine carboxamide.

## References

(1) Meldal, M.; Schoffelen, S. Recent Advances in Covalent, Site-Specific Protein Immobilization. F1000Research 2016, 5, 2303. https://doi.org/10.12688/f1000research.9002.1.

(2) Romero-Fernández, M.; Paradisi, F. Protein Immobilization Technology for Flow Biocatalysis. Curr. Opin. Chem. Biol. 2020, 55, 1–8. https://doi.org/10.1016/j.cbpa.2019.11.008.

(3) Steen Redeker, E.; Ta, D. T.; Cortens, D.; Billen, B.; Guedens, W.; Adriaensens, P. Protein Engineering for Directed Immobilization. Bioconjug. Chem. 2013, 24 (11), 1761–1777. https://doi.org/10.1021/bc4002823.

(4) Goding, J. W. Use of Staphylococcal Protein A as an Immunological Reagent. J. Immunol. Methods 1978, 20, 241–253. https://doi.org/10.1016/0022-1759(78)90259-4.

(5) Pelton, R. Bioactive Paper - A Paper Science Perspective. In Advances in Pulp and Paper Research, Oxford 2009 (S.J.I’Anson, ed.); FRC, Manchester, 2018; pp 1095–1145.

(6) Manson, L. A.; Verastegui-Cerdan, E.; Sporer, R. A Quantitative Disc Radioimmunoassay for Antibodies Directed against Membrane-Associated Antigens. Curr. Top. Microbiol. Immunol. 1978, 81, 232–234.

(7) Beddows, C. G.; Mirauer, R. A.; Guthrie, J. T. Immobilization of β-Galactosidase and Other Enzymes onto p-Amino-Carbanilated Cellulose Derivatives. Biotechnol. Bioeng. 1980, 22 (2), 311–321. https://doi.org/10.1002/bit.260220206.

(8) Hong, W.; Jeong, S.-G.; Shim, G.; Kim, D. Y.; Pack, S. P.; Lee, C.-S. Improvement in the Reproducibility of a Paper-Based Analytical Device (PAD) Using Stable Covalent Binding between Proteins and Cellulose Paper. Biotechnol. Bioprocess Eng. BBE 2018, 23 (6), 686–692. https://doi.org/10.1007/s12257-018-0430-2.

(9) Tomme, P.; Boraston, A.; McLean, B.; Kormos, J.; Creagh, A. L.; Sturch, K.; Gilkes, N. R.; Haynes, C. A.; Warren, R. A. J.; Kilburn, D. G. Characterization and Affinity Applications of Cellulose-Binding Domains. J. Chromatogr. B. Biomed. Sci. App. 1998, 715 (1), 283–296. https://doi.org/10.1016/S0378-4347(98)00053-X.

(10) Boraston, A. B.; Bolam, D. N.; Gilbert, H. J.; Davies, G. J. Carbohydrate-Binding Modules: Fine-Tuning Polysaccharide Recognition. Biochem. J. 2004, 382 (Pt 3), 769–781. https://doi.org/10.1042/BJ20040892.

(11) Sidar, A.; Albuquerque, E. D.; Voshol, G. P.; Ram, A. F. J.; Vijgenboom, E.; Punt, P. J. Carbohydrate Binding Modules: Diversity of Domain Architecture in Amylases and Cellulases From Filamentous Microorganisms. Front. Bioeng. Biotechnol. 2020, 8, 871. https://doi.org/10.3389/fbioe.2020.00871.

(12) Attia, M. A.; Brumer, H. New Family of Carbohydrate-Binding Modules Defined by a Galactosyl-Binding Protein Module from a Cellvibrio Japonicus Endo-Xyloglucanase. Appl. Environ. Microbiol. 2021, 87 (5), e02634–20. https://doi.org/10.1128/AEM.02634-20.

(13) Ong, E.; Gilkes, N. R.; Miller, R. C.; Antony, R.; Warren, J.; Kilburn, D. G. Enzyme Immobilization Using a Cellulose-Binding Domain: Properties of a β-Glucosidase Fusion Protein. Enzyme Microb. Technol. 1991, 13 (1), 59–65. https://doi.org/10.1016/0141-0229(91)90189-H.

(14) Rodriguez, B.; Kavoosi, M.; Koska, J.; Creagh, A. L.; Kilburn, D. G.; Haynes, C. A. Inexpensive and Generic Affinity Purification of Recombinant Proteins Using a Family 2a CBM Fusion Tag. Biotechnol. Prog. 2004, 20 (5), 1479–1489. https://doi.org/10.1021/bp0341904.

(15) Hong, J.; Wang, Y.; Ye, X.; Zhang, Y.-H. P. Simple Protein Purification through Affinity Adsorption on Regenerated Amorphous Cellulose Followed by Intein Self-Cleavage. J. Chromatogr. A 2008, 1194 (2), 150–154. https://doi.org/10.1016/j.chroma.2008.04.048.

(16) Sugimoto, N.; Igarashi, K.; Samejima, M. Cellulose Affinity Purification of Fusion Proteins Tagged with Fungal Family 1 Cellulose-Binding Domain. Protein Expr. Purif. 2012, 82 (2), 290– 296. https://doi.org/10.1016/j.pep.2012.01.007.

(17) Carrick, B. H.; Hao, Linxuan; Smaldino, P. J.; Engelke, D. R. A Novel Recombinant DNA System for High Efficiency Affinity Purification of Proteins in Saccharomyces Cerevisiae. G3 Bethesda 6, 573–578.

(18) Morag, E.; Lapidot, A.; Govorko, D.; Lamed, R.; Wilchek, M.; Bayer, E. A.; Shoham, Y. Expression, Purification, and Characterization of the Cellulose-Binding Domain of the Scaffoldin Subunit from the Cellulosome of Clostridium Thermocellum. Appl. Environ. Microbiol. 1995, 61 (5), 1980–1986. https://doi.org/10.1128/AEM.61.5.1980-1986.1995.

(19) Studier, F. W.; Moffatt, B. A. Use of Bacteriophage T7 RNA Polymerase to Direct Selective High-Level Expression of Cloned Genes. J. Mol. Biol. 1986, 189 (1), 113–130. https://doi.org/10.1016/0022-2836(86)90385-2.

(20) Leung, K. K. K.; Shilton, B. H. Chloroquine Binding Reveals Flavin Redox Switch Function of Quinone Reductase 2. J. Biol. Chem. 2013, 288 (16), 11242–11251. https://doi.org/10.1074/jbc.M113.457002.

(21) Leung, K. K. K.; Shilton, B. H. Binding of DNA-Intercalating Agents to Oxidized and Reduced Quinone Reductase 2. Biochemistry 2015, 54 (51), 7438–7448. https://doi.org/10.1021/acs.biochem.5b00884.

(22) Tropea, J. E.; Cherry, S.; Waugh, D. S. Expression and Purification of Soluble His(6)-Tagged TEV Protease. Methods Mol. Biol. Clifton NJ 2009, 498, 297–307. https://doi.org/10.1007/978-1-59745-196-3_19.

(23) Peterson, G. L. Determination of Total Protein. Methods Enzymol. 1983, 91, 95–119. https://doi.org/10.1016/s0076-6879(83)91014-5.

(24) Laemmli, U. K. Cleavage of Structural Proteins during the Assembly of the Head of Bacteriophage T4. Nature 1970, 227 (5259), 680–685. https://doi.org/10.1038/227680a0.

(25) Kuzmič, P. Program DYNAFIT for the Analysis of Enzyme Kinetic Data: Application to HIV Proteinase. Anal. Biochem. 1996, 237 (2), 260–273. https://doi.org/10.1006/abio.1996.0238.

(26) Zhao, Q.; Yang, X. L.; Holtzclaw, W. D.; Talalay, P. Unexpected Genetic and Structural Relationships of a Long-Forgotten Flavoenzyme to NAD(P)H:Quinone Reductase (DT-Diaphorase). Proc. Natl. Acad. Sci. 1997, 94 (5), 1669–1674. https://doi.org/10.1073/pnas.94.5.1669.

(27) Islam, F.; Leung, K.; Walker, M. D.; Al Massri, S.; Shilton, B.H. The Unusual Cosubstrate Specificity of NQO2: Conservation Throughout the Amniotes and Implications for Cellular Function. Front. Pharmacol. 2022, in revision.

(28) Vella, F.; Ferry, G.; Delagrange, P.; Boutin, J. A. NRH:Quinone Reductase 2: An Enzyme of Surprises and Mysteries. Biochem. Pharmacol. 2005, 71 (1–2), 1–12. https://doi.org/10.1016/j.bcp.2005.09.019.

(29) Tormo, J.; Lamed, R.; Chirino, A. J.; Morag, E.; Bayer, E. A.; Shoham, Y.; Steitz, T. A. Crystal Structure of a Bacterial Family-III Cellulose-Binding Domain: A General Mechanism for Attachment to Cellulose. EMBO J. 1996, 15 (21), 5739–5751. https://doi.org/10.1002/j.1460-2075.1996.tb00960.x.

(30) Shpigel, E.; Goldlust, A.; Eshel, A.; Ber, I. K.; Efroni, G.; Singer, Y.; Levy, I.; Dekel, M.; Shoseyov, O. Expression, Purification and Applications of Staphylococcal Protein A Fused to Cellulose-Binding Domain. Biotechnol. Appl. Biochem. 2000, 31 (3), 197. https://doi.org/10.1042/BA20000002.

(31) Cao, Y.; Zhang, Q.; Wang, C.; Zhu, Y.; Bai, G. Preparation of Novel Immunomagnetic Cellulose Microspheres via Cellulose Binding Domain-Protein A Linkage and Its Use for the Isolation of Interferon α-2b. J. Chromatogr. A 2007, 1149 (2), 228–235. https://doi.org/10.1016/j.chroma.2007.03.032.

(32) Harlow, Ed; Lane, David. Antibodies A Laboratory Manual; Cold Srping Harbor Laboratory, 1988.

(33) Natarajan, S.; Jayaraj, J.; Prazeres, D. M. F. A Cellulose Paper-Based Fluorescent Lateral Flow Immunoassay for the Quantitative Detection of Cardiac Troponin I. Biosensors 2021, 11 (2), 49. https://doi.org/10.3390/bios11020049.

(34) Berdichevsky, Y.; Lamed, R.; Frenkel, D.; Gophna, U.; Bayer, E. A.; Yaron, S.; Shoham, Y.; Benhar, I. Matrix-Assisted Refolding of Single-Chain Fv– Cellulose Binding Domain Fusion Proteins. Protein Expr. Purif. 1999, 17 (2), 249–259. https://doi.org/10.1006/prep.1999.1125.

(35) Shpigel, E.; Elias, D.; Cohen, I. R.; Shoseyov, O. Production and Purification of a Recombinant Human Hsp60 Epitope Using the Cellulose-Binding Domain InEscherichia Coli. Protein Expr. Purif. 1998, 14 (2), 185–191. https://doi.org/10.1006/prep.1998.0929.

(36) Ramos, R.; Domingues, L.; Gama, M. Escherichia Coli Expression and Purification of LL37 Fused to a Family III Carbohydrate-Binding Module from Clostridium Thermocellum. Protein Expr. Purif. 2010, 71 (1), 1–7. https://doi.org/10.1016/j.pep.2009.10.016.

(37) Ramos, R.; Moreira, S.; Rodrigues, A.; Gama, M.; Domingues, L. Recombinant Expression and Purification of the Antimicrobial Peptide Magainin-2. Biotechnol. Prog. 2013, 29 (1), 17–22. https://doi.org/10.1002/btpr.1650.

(38) Miller, E. A.; Baniya, S.; Osorio, D.; Al Maalouf, Y. J.; Sikes, H. D. Paper-Based Diagnostics in the Antigen-Depletion Regime: High-Density Immobilization of RcSso7d-Cellulose-Binding Domain Fusion Proteins for Efficient Target Capture. Biosens. Bioelectron. 2018, 102, 456–463. https://doi.org/10.1016/j.bios.2017.11.050.

